# Brunner Syndrome associated MAOA dysfunction in human induced dopaminergic neurons results in dysregulated NMDAR expression and increased network activity

**DOI:** 10.1101/741108

**Authors:** Y. Shi, J.R. van Rhijn, M. Bormann, B. Mossink, M. Frega, M. Hakobjan, S. Kittel-Schneider, D. Schubert, H. Brunner, B. Franke, N. Nadif Kasri

## Abstract

Monoamine oxidase A (MAOA) is an enzyme that catalyzes the degradation of dopamine, noradrenaline, and serotonin. Regulation of monoamine neurotransmitter abundance through MAOA activity strongly affects motor control, emotion, and cognitive function. Mutations in MAOA cause Brunner Syndrome, which is characterized by impulsive aggressive behavior and mild intellectual disability (ID). The impaired MAOA activity in Brunner Syndrome patients results in bioamine aberration, but it is currently unknown how this affects neuronal function. MAOA is highly expressed in serotonergic and dopaminergic neurons, and dysfunction of both neurotransmission systems is associated with aggressive behavior in mice and humans. Research has so far mainly focused on the serotonergic system. Here, we generated human induced pluripotent stem cell-derived induced dopaminergic neurons (iDANs) from individuals with known MAOA mutations, to investigate MAOA-dependent effects on dopamine neuronal function in the context of Brunner Syndrome. We assessed iDAN lines from three patients and combined data from morphological analysis, gene expression, single-cell electrophysiology, and network analysis using micro-electrode arrays (MEAs). We observed mutation-dependent functional effects as well as overlapping changes in iDAN morphology. The most striking effect was a clear increase in N-methyl-D-aspartate (NMDA) receptor mRNA expression in all patient lines. A marked increase was also seen in coordinated network activity (network bursts) on the MEA in all patient lines, while single-cell intrinsic properties and spontaneous excitatory postsynaptic currents activity appeared normal. Together, our data indicate that dysfunction of MAOA leads to increased coordinated network activity in iDANs, possibly caused by increased synaptic NMDA receptor expression.

## Introduction

Monoamines, such as dopamine and serotonin, have an important role in brain function, and dysregulation of monoaminergic pathways is associated with several mental disorders including schizophrenia, autism spectrum disorder, and attention deficit/hyperactivity disorder (ADHD)^1^. The abundance of monoamines is tightly regulated by monoamine oxidases (MAOs), which catabolize the monoaminergic neurotransmitters^2^. Disruption of MAO activity can thus have profound consequences on normal brain function^3^. Brunner Syndrome is a neurodevelopmental disorder characterized by hemizygous mutation of the X-linked monoamine oxidase, and was first described in a large Dutch kindred with non-dysmorphic borderline intellectual disability (ID) and prominent impulsive aggressive behavior^4,5^. Recently, three more families have been reported in which affected individuals have Brunner Syndrome and carry either nonsense or missense mutations of *MAOA*^6,7^, which strengthens the association between MAOA dysfunction and Brunner Syndrome.

Changes in monoaminergic regulation, especially of serotonin and dopamine, have also been associated with aggressive behavior in animal models, both in wild-type animals^8–11^ as well as in genetic models for neurodevelopmental disorders^12^. Transgenic mouse models of *Maoa* have allowed the exploration of mechanistic associations between monoamine regulation and aggressive behavior. Hemizygous *Maoa* mutant mice show abnormally high levels of aggressive behavior and disturbed monoamine metabolism, similar to the patient situation^13–15^. *Maoa*-deficient mice also display alterations in brain development, with aberrant organization of the primary somatosensory cortex^13^ and increased dendritic arborization of pyramidal neurons in the orbitofrontal cortex^15^. Furthermore, *Maoa* has been implicated in the regulation of synaptic neurotransmitter receptors, as *Maoa* knockout mice show increased N-methyl-D-aspartate (NMDA) receptor subunit expression in the prefrontal cortex^16^. Taken together, these data suggest that *Maoa* can influence both structural and functional aspects of neurodevelopment.

MAOA is expressed in different neuronal as well as glial cell types in the brain^*17*^. However, not all of these cell types are thought to be associated with the behavioral phenotypes observed in *MAOA* mutation carriers. So far, particular emphasis has been given to the serotonergic system in MAOA research, but MAOA is also abundantly expressed in dopaminergic neurons^18,19^, and aggressive behavior is highly associated with aberrant dopamine pathway function^1,20^. The current advances in the generation of human pluripotent stem cell (iPSC)-induced neurons (iNeurons) enable us to generate a homogenous culture of a single neuronal cell type. We therefore generated iPSC-derived induced dopaminergic neurons (iDANs), which offer a unique opportunity to investigate the cellular and molecular mechanisms specifically affected by MAOA dysfunction in dopaminergic neurons. We generated iDANs from three independent male individuals with unique *MAOA* mutations and from two independent male controls. Combining data from morphological analysis, gene expression, single-cell electrophysiology, and network analysis using micro electrode arrays (MEAs), we suggest that alterations in dopaminergic neuron development and function may also be relevant to behavioural and cognitive dysfunction in Brunner Syndrome.

## Results

### Generation of control and Brunner Syndrome patient-derived iDANs

Using a protocol adapted from^21^, we successfully differentiated iPSCs into iDANs after 55 days *in vitro* (DIV55, **Figure 1a**). We generated iDANs from two independent control (control-1 and control-2) and three independent patient iPSC lines (Patient-NE8, c.886C>T, p.Q296*; Patient-ME8, c.797_798delinsTT, p.C266F; Patient-ME2, c.133C>T, p.R45W; **Figure 1b**, for extended information see **Supplementary Table S2**). Neuronal identity was confirmed by microtubule-associated protein 2 (MAP2) expression and iDAN identity by expression of tyrosine hydroxylase (TH) (**Figure 1c**). Cell type specificity was comparable between lines, with more than 90% of MAP2-expressing neurons staining positive for TH (**Figure 1d**). Interestingly, mRNA levels of MAOA were only altered compared to the healthy controls in the Patient-NE8 line, which has a nonsense mutation (**Figure 1e**). The observed reduction might have been caused by nonsense-mediated mRNA decay, as shown earlier in human fibroblasts with the same mutation^25^. We used these five lines in subsequent experiments to investigate how MAOA dysfunction affects iDAN development and function.

**Figure 1:**
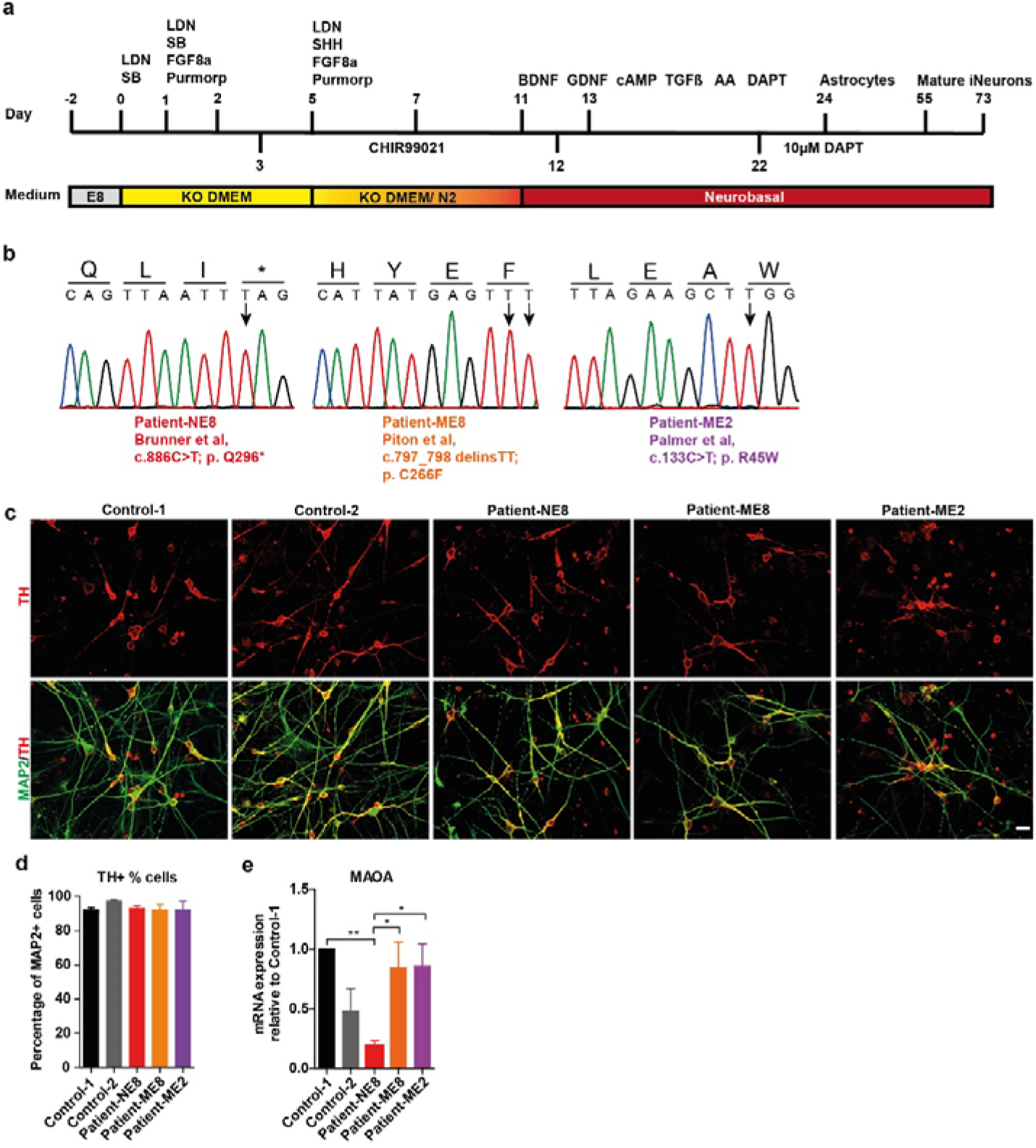
Differentiation of dopaminergic neurons derived from human iPSCs. (a) Schematic overview the protocol used to generate dopaminergic neurons from human iPSCs. (b) The sanger sequencing used to validate the *MAOA* mutations present in iPSCs lines used in this study. Patient-NE8 carries a nonsense mutation at exon 8, Patient-ME2 and Patient-ME8 carry missense mutations in exon 2 and exon 8, respectively. (c) Representative images of DIV55 dopaminergic iDANs labeled by TH (red) and MAP2 (green) (Scale bar=20 μm). (d) The percentage of TH-positive neurons (among MAP2-positive cells) did not differ between cell lines (n=3). (e) MAOA expression in DIV73 iDANs as quantified by qPCR. MAOA mRNA expression is only significantly reduced in the patient-NE8 cell line compared to control 1 or the other patient cell lines. However, expression between patient-NE8 and control-2 was comparable. Significance was determined by one-way ANOVA followed by post-hoc Tukey’s multiple comparison(n≥5). *p<0.05; **p<0.01.

### Mutation-specific effects of *MAOA*-disruption on iDAN morphology

Dendritic properties of neurons in the orbitofrontal cortex are known to be altered in *Maoa* hemizygous knockout mice^15^. To assess whether MAOA dysfunction also affects the morphology of iDANs, we compared dendritic properties and soma size of control and patient iDANs by quantitative morphometric analysis of the somatodendritic compartment. ME8, NE8 and ME2 patient lines revealed mutation-dependent effects on iDAN morphology. Only iDANs with ME8 mutation showed increases in soma size (**Figure 2b**) and dendrite complexity (**Figure 2c,d,e,f**). Additionally, iDANs with the ME8 and ME2 mutations showed a significant increase in total covered surface (Convex Hull analysis, **Figure 2g**), whilst iDANs with the NE8 mutation showed no differences in neuronal complexity. Next, we investigated if these differences in morphology could affect the number of excitatory synapses present on iDANs, since changes in neuronal complexity^26,27^ and synapse density^28^ are widely associated with neurodevelopmental disorders. Indeed, synapse density was significantly decreased in iDANs from all patients compared to those from control-1 (**Figure 2h, i**). This suggests that differences in synapse density might be caused by a molecular mechanism affected across mutations, whereas the drastic difference in iDAN dendritic morphology might be mutation-specific.

**Figure 2:**
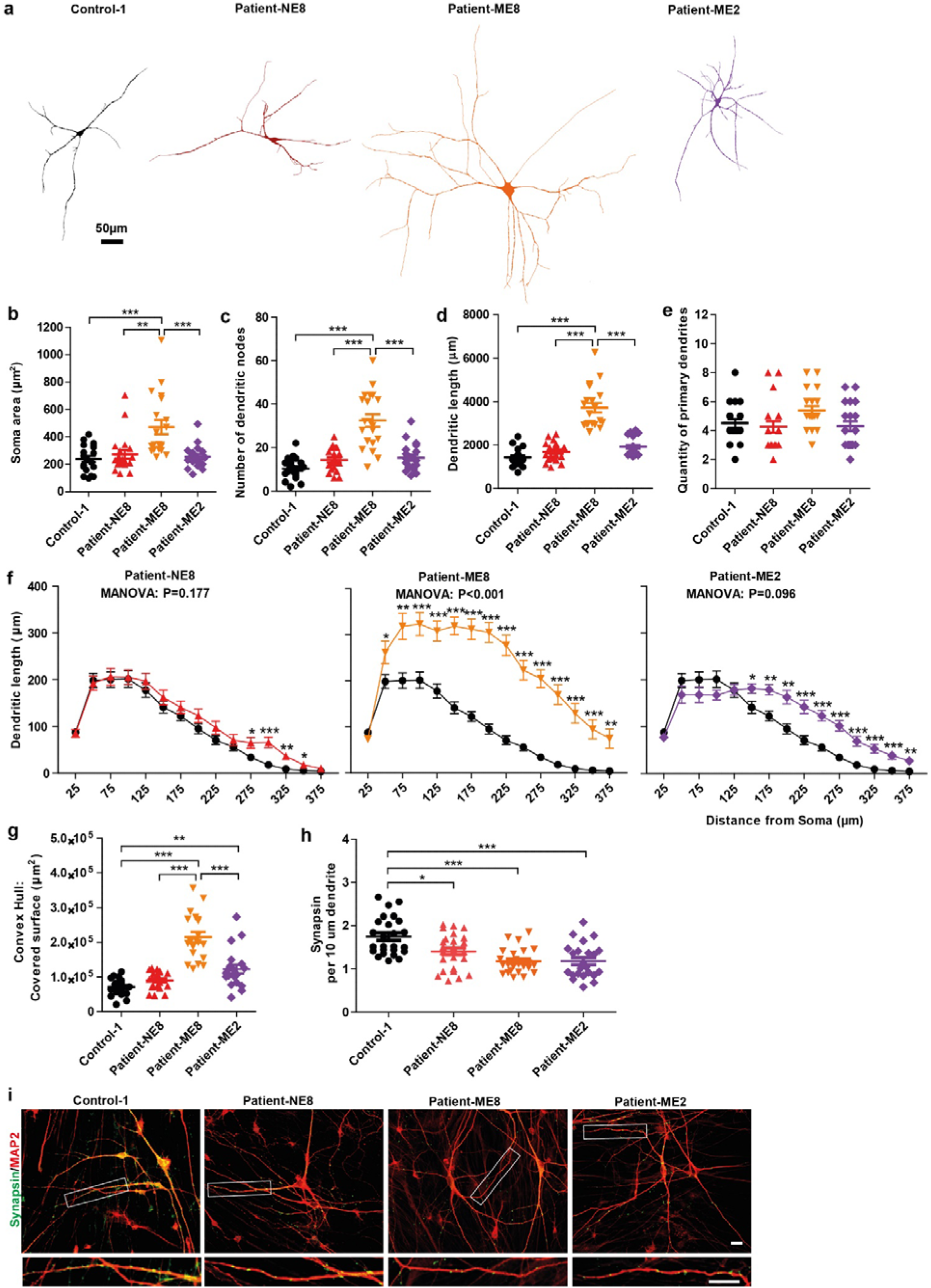
Morphological comparison of control and patient derived iDANs. (a) Representative images of reconstructed iDANs (DIV73) iDANs (N/n = 3/20). The quantitative parameters of neuronal morphology of the day 73 iDANs were compared with the control-1 by multivariate analysis of variance (MANOVA) in terms of soma area (b), number of dendritic nodes (c), dendritic length (d), quantity of primary dendrites (e) and Convex Hull: covered surface (g) followed by Bonferroni post-hoc correction. (f) Sholl analysis of dendritic length of DIV73 iDANs at different distances measured from soma center. Statistical analysis between groups was done using MANOVA with Wilks’ lambda test. Synapsin density was decreased in the *MAOA*-mutant iDANs when compared with Control-1 by One-way ANOVA followed with Bonferroni post-hoc correction(h) (n≥23 per genotype). (i) Representative images of control and patient iDANs. Dendrites are labeled with MAP2 (red) and synapses with synapsin (green), scale bar = 20μm. Inset shows a single stretch of dendrite with clearly visible synapses, scale bar = 10um. All data is represented as mean ±SEM, asterisks indicate significant differences between genotypes. *p<0.05; **p<0.01; ***p<0.001. N/n = number of batches / number of neurons.

### MAOA dysfunction leads to increased NMDAR expression in iDANs

The alterations in morphology and synapse density in *MAOA* hemizygous mutation iDANs prompted us to investigate whether gene expression of synapse-specific genes might be affected by MAOA dysfunction. In excitatory neurons, both α-amino-3-hydroxy-5-methyl-4-isoxazolepropionic acid receptors (AMPARs) and N-methyl D-aspartate receptors (NMDARs) are especially relevant for glutamatergic synaptic neurotransmission. Both AMPARs and NMDARs are heteromeric ion channels comprised of four subunits, and the ion channel composition conveys differences in ion selectivity as well as opening and closing kinetics. We therefore investigated the expression of the four main isoforms of the AMPA receptor subunits GluA1, 2, 3 and 4 (encoded by *GRIA1, 2, 3*, and *4*, respectively) as well as the expression of the NR1, NR2A, and NR2B subunits (encoded by *GRIN1, 2A* and *2B*, respectively) of the NMDA receptor.

When we compared control iDANs at DIV73 to our patient-derived iDANs, no difference in AMPAR subunit expression was seen (**Figure S3a-d**). This suggests AMPAR-mediated neurotransmission is not affected in dopaminergic neurons by mutations in *MAOA*. Following evidence from *Maoa* knockout mice showing that NMDAR expression is altered in the prefrontal cortex^16^, we also investigated NMDAR in our patient-derived iDANs compared to controls. No difference in gene expression was found for *GRIN1* (**Figure 3a**), but increased expression was seen for both *GRIN2A* and *GRIN2B* in the patient-derived iDANs (**Figure 3b,c**), similar to the earlier observations in *Maoa* knockout mice^16^. Together, this data suggests that excitatory neurotransmitter receptor expression at the synapse is altered by MAOA dysfunction.

**Figure 3:**
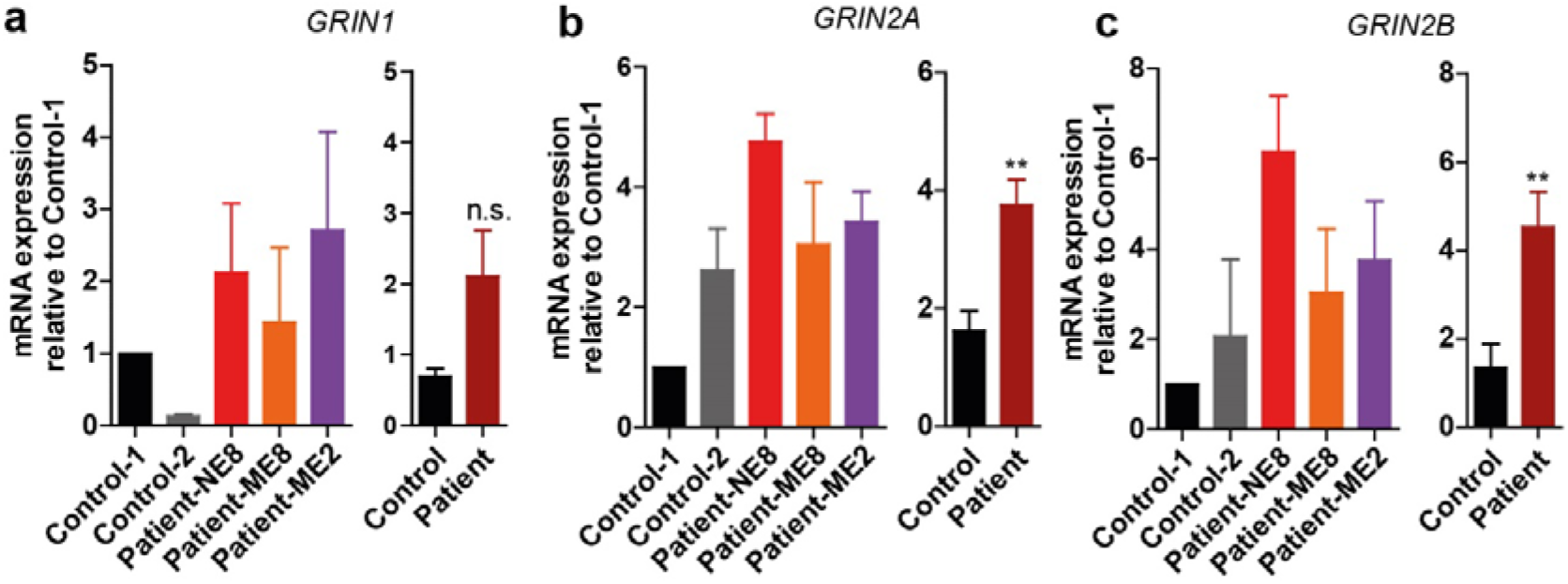
Patient-derived iDANs exhibit higher expression of NMDA receptor subunits *GRIN2A* and *GRIN2B* mRNA. The expression of genes encoding NMDA receptor subunits *GRIN1*(a), *GRIN2A*(b) and *GRIN2B*(c) was analyzed by qPCR in DIV73 iDANs. The mRNA expression was normalized to the mean of Control-1. Statistical analysis was done with Mann-Whitney U-test between the group of control and patient. Data are mean ±SEM, asterisks indicate significant differences between groups. **p<0.01. (n≥ 5). N.s.: not significant.

### Intrinsic properties and AMPAR-mediated excitatory activity are not affected by MAOA mutation

As NMDAR expression is tightly linked to burst firing in dopaminergic neurons^29^ and since differences in neuronal morphology and synapse density, as we observed them, can have drastic consequences for neuronal activity, we next explored if physiological activity is altered by MAOA dysfunction in iDANs. First, we explored whether MAOA dysfunction affects synapse physiology through altered excitatory neuron function using whole-cell patch clamp. Comparing the intrinsic properties of the control lines, we found no significant differences between control-1 and control-2 (**Figure 4a-d**) and pooled the data for comparison with the patient-derived lines. No differences in intrinsic properties were observed between the pooled data of the control lines and the three patient lines (**Figure 4e-h**). To assess whether AMPAR-mediated inputs are affected by MAOA dysfunction, we measured spontaneous excitatory postsynaptic currents (sEPSCs). Again, we first compared control lines and found no differences in sEPSC amplitude or frequency (**Figure 4i-k**). Comparison of the pooled control data to data from the individual patients revealed that AMPAR-mediated excitatory activity was not affected by mutations in *MAOA* (**Figure 4i,l,m**); these findings are consistent with the absence of differences in the expression of GluA subunit mRNA between patients and controls.

**Figure 4:**
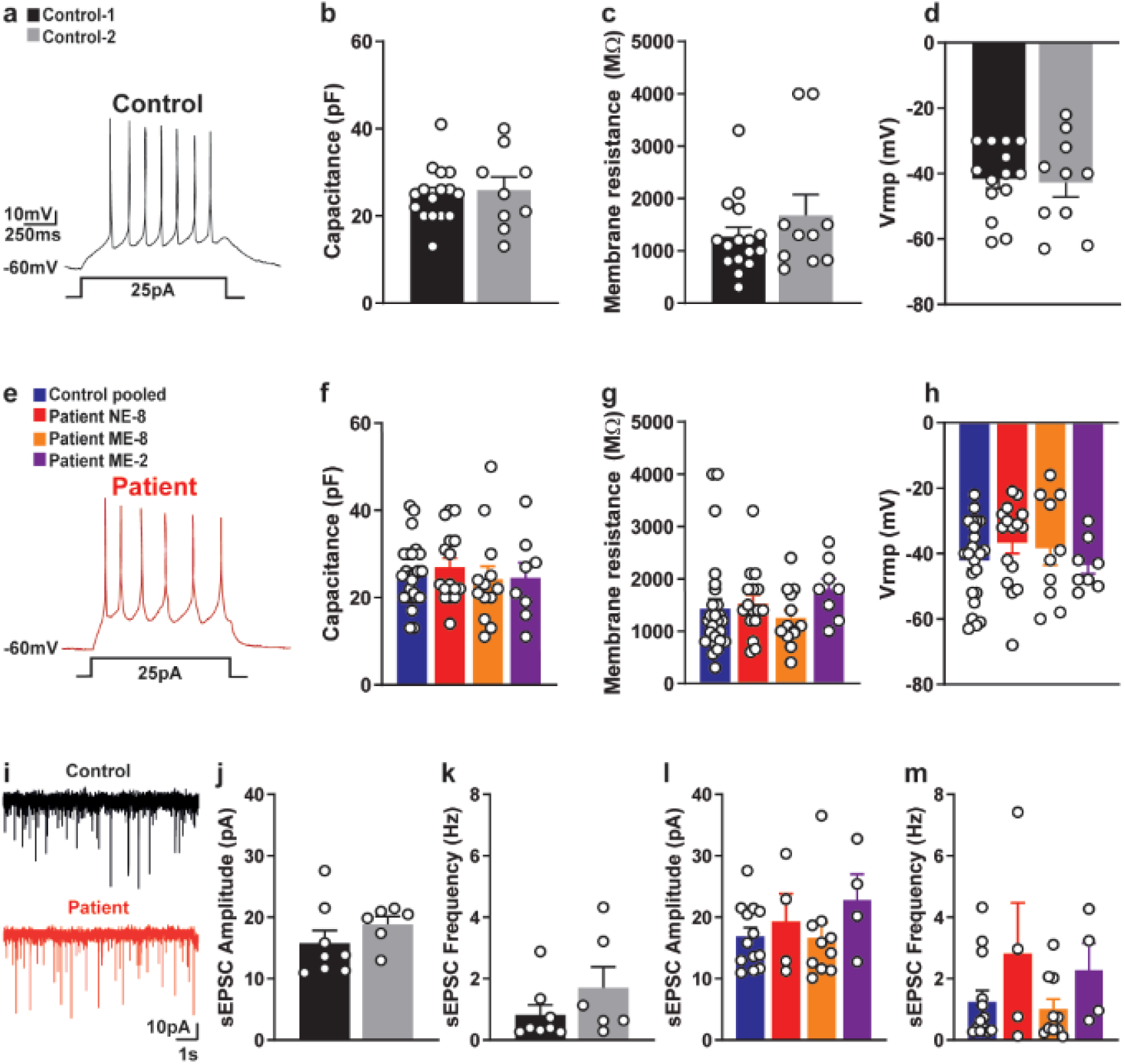
MAOA dysfunction does not affect intrinsic properties or sEPSC activity. (a) Representative trace of action potentials generated by DIV55 control-1 iDANs. No significant differences were observed for capacitance (b), membrane resistance (c) or resting membrane potential (d) between control-1 and control-2. (e) Representative trace of action potentials generated by DIV55 NE-8 iDANs. No differences were found between pooled control data and the patient lines, nor between the patient lines separately, in capacitance (f) membrane resistance (g) or resting membrane potential (h). (i) Representative traces of sEPSC activity in DIV73 control-1 and patient-NE8 derived iDANs. sEPSC amplitude (j) and frequency (k) were similar between control-1 and control 2, as well as between pooled control data and patient lines (l,m). Data was analyzed by Student’s T-test (control-1 vs control-2) or ANOVA (control vs patient lines) with correction for multiple testing. Data are shown as mean ± SEM.

### MAOA dysfunction leads to increased iDAN network activity recorded by MEA

Since we found no differences in intrinsic properties and spontaneous AMPAR-driven activity at the single cell level, we next explored whether the increased NMDAR mRNA expression in *MAOA*-mutant iDAN lines leads to changes in neuronal activity at the network level. Therefore, we measured network activity of iDANs with MEA. Spontaneous random activity was clearly visible for all five iDAN cell lines at DIV73 (**Figure 5a**), and robustly higher mean firing rates were seen for all patient lines compared to the activity in both control lines (**Figure 5b,c**). Furthermore, whilst the control lines exhibited almost no coordinated network bursts, these were clearly visible in each of the patient-derived lines (**Figure 5a**, colored blocks in traces), and their frequency was significantly higher (**Figure 5d,e**). This suggests that both missense and nonsense mutations of *MAOA* lead to increased activity in excitatory dopaminergic neurons, possibly through upregulation of NMDAR-driven network excitation.

**Figure 5:**
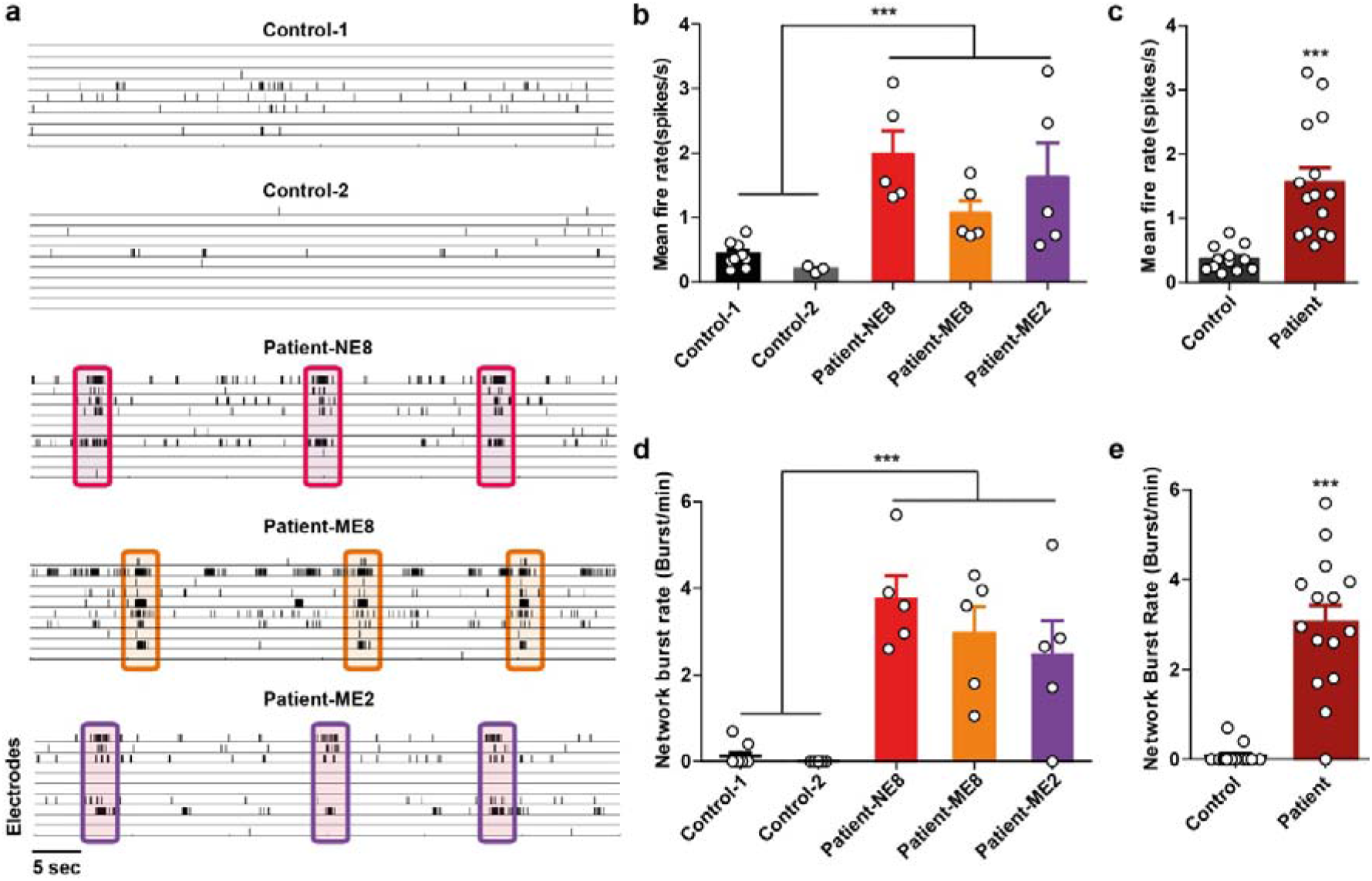
Increased neuronal network activity in the MAOA mutant iDANs. (a) Representative rastplot of the action potential activity across 9 micro-electrodes of day 73 iDANs. Each detected action potential is indicated as a black bar. Neuronal network burst events are highlighted by boxes in the *MAOA*-mutant iDANs. Both the mean fire rate, which is the average action potential frequency detected by an active electrode (b,c), and the network burst rate (d,e) were increased in *MAOA*-mutant iDANs. Statistical analysis was performed with Mann Whitney U-test (n≥5 per genotype). Data are mean ±SEM; asterisks indicate significant differences between controls and *MAOA*-mutant iDANs. ***p<0.001.

## Discussion

Dysfunction of MAOA in humans results in Brunner Syndrome. Here, we investigated Brunner Syndrome in a human iPSC-derived neuronal model for the first time, choosing dopaminergic neurons as our model. We generated two control and three patient lines and compared morphology, expression of synaptic genes, and neuronal physiology at the single-cell and network level. Confirming findings obtained in a *Maoa* knockout mouse model^15,16^, we found neuronal complexity to be higher in dopaminergic neurons derived from humans carrying *MAOA* mutations. Our work further extends current knowledge by showing that such differences are mutation-dependent, as the p.C266F mutation (ME8) showed the strongest differences in iDAN morphology, whereas in the p.Q296* (NE8) line such alterations were less severe, and no differences were seen in the p.R45W (ME2) line. This suggests that different mutations might uniquely affect the function of the MAOA, as has been hypothesized in *in vitro* studies of MAOA function^30^. In addition to the mutation-specific effects on neuronal morphology, we observed several differences in neuronal function shared amongst all mutations. Firstly, we observed an increase in NMDAR mRNA expression, whilst AMPAR mRNA levels were unchanged. While intrinsic properties and spontaneous activity of single cells were unaffected, neuronal activity at the network level was robustly higher in all patient-derived lines compared to control iDANs.

The differences in spontaneous activity and in NMDAR expression could be related. Firstly, it has been shown that in *Maoa* knockout mice NMDAR expression is increased, which affects NMDAR-dependent currents in prefrontal cortex *ex vivo* slice preparations of these mice^16^. Furthermore, increased NMDAR-expression is able to enhance spontaneous neuronal activity during development^31^, and dysfunctional enhancement of NMDAR signaling has been described in other neurodevelopmental disorders^32^. However, how *MAOA* mutation and increased NMDAR expression are related on a mechanistic level needs to be further explored.

Synapse density was found reduced across patient-derived lines. This could be seen a compensatory mechanism in the case of the higher neuronal complexity in the ME8 patient line, but this cannot explain the reduced synapse density in the other patient lines. The expression and localization of NMDARs has been shown to regulate dendritic arborization^33–35^. It is therefore conceivable that the increased dendrite complexity and the changes in synapse density in the *MAOA*-mutant lines can be partially explained by the increased NMDAR expression.

The individuals we investigated in our study all carry rare mutations, which lead to either complete loss or reduced activity of the MAOA enzyme. The MAOA promoter also can exhibit different levels of transcriptional activity^39^ through a common variable number of tandem repeats (VNTR) polymorphism in the *MAOA* regulatory region, located 1.2kb upstream of the transcription initiation site. Low activity alleles of *MAOA* that show similar transcriptional activity as the missense mutations present in Brunner Syndrome patients have been associated with the anti-social behavior in individuals subjected to childhood maltreatment^40,41^. Furthermore, an association has been observed between the severity of ASD features and the presence of *MAOA* low activity alleles^42,43^, which suggests that *MAOA* also plays a role in ASD pathology^38^. Lastly, impulsive aggressive behavior has been described as a feature of other, genetically complex neurodevelopmental disorders such as ADHD^44^ and ASD^45^, in which monoaminergic signaling seems to be involved but the association with *MAOA* allele activity has not yet been extensively explored. Interestingly, using magnetic resonance imaging (MRI) in a large group of healthy individuals, we recently showed differences in structural and functional network architecture between carriers of low and high activity MAOA alleles, where those with low activity alleles had more extensive networks. Investigation of MAOA function in neurons derived from individuals with low or high activity MAOA alleles might thus help us further understand the molecular mechanisms regulated by MAOA and how MAOA activity might influence impulsive aggressive behavior in the general population.

Lastly, MAOA dysfunction does not exclusively affect the dopaminergic system, as it also results in increased serotonin levels in both human and mice^5,6,7,13,15,46^. Additionally, the serotonergic pathway is, similar to the dopaminergic pathway, associated with aggression and impulsivity^47^. Previously, it was suggested the increased serotonin levels are an indirect effect of reduced MAOA activity, as MAOA was shown not to be expressed in serotonergic neurons. However, recent single cell RNAseq profiling revealed MAOA is expressed both in neural progenitor cells and monoaminergic neurons, including both dopaminergic and serotonergic neurons^19,48^.

This study should be viewed in the context of its strengths and limitations. This is the first study on Brunner Syndrome investigating the molecular, cellular, and circuit alterations underlying the phenotype of the patients in iPSC-derived homogeneous neuronal cultures. The wide range of methods used and the integration of findings allowed us to validate results of animal models and extend our insight into the biological underpinnings of Brunner Syndrome further.

In conclusion, we established iDANs as a pure iPSC-derived human neuron culture that can be used to investigate disorders, which specifically affect monoaminergic pathways. Our data on MAOA dysfunction using this tool suggest that dopaminergic neurons may also be altered in humans carrying *MAOA* mutations, extending research focused on the serotonergic system. We confirm work performed in rodent models and non-neuronal transgenic models and extend such knowledge further by identifying both mutation-specific as well as general effects of MAOA dysfunction. Our findings of altered network activity may also link to human MRI studies showing altered structural and functional networks in carriers of a functional polymorphism in the *MAOA* gene. Beyond investigation of the biological underpinnings of neurodevelopmental disorders like Brunner Syndrome, the infrastructure we built may also provide a platform to investigate potential (individualized) therapeutic interventions for disorders in which monoamine regulation is disrupted.

## Materials and Methods

### Cell Culture of iPSCs

The iPSC line control-1 was derived from a dermal fibroblast biopsy of a male healthy volunteer. iPSC line control-2 was obtained from^50^ and was generated from a healthy male donor using an episomal reprogramming^51^. The iPSC lines patient-NE8, patient-ME8 and patient-ME2 that carry *MAOA* mutations were derived from dermal fibroblast biopsies of male patients described previously^4–7^. The fibroblasts were reprogrammed by introducing four Yamanaka factors: Sox2, Klf4, Oct4, c-Myc via a lentivirus-based method. All iPSC reprogramming was done by the Radboudumc Stem Cell Technology Center (SCTC). iPSC pluripotency was characterized by immunocytochemistry and qPCR (supplemental figure S1). The iPSCs were maintained in Essential 8 complete medium with 100 μg/ml penicillin/streptomycin or 100 μg/ml Primocin (InvivoGen) on vitronectin (VTN-N)-coated cell culture plates (Corning) at 37°C/5% CO2. Cell colonies were passaged by 0.5 mM ultrapure EDTA treatment for 2 or 3 minutes in DPBS without calcium and magnesium. Unless indicated otherwise, all the reagents were bought from Thermo Fisher Scientific Inc.

### Differentiation of iPSCs into dopaminergic neurons

The iPSC colonies were split into single cells by Accutase treatment for 5 min at 37°C two days before starting the differentiation, and 1-2x 10^4^/cm^2^ cells were replated on a vitronectin-coated plate in Essential 8 complete medium supplemented with either 2uM thiazovivin(Sigma) or 1X RevitaCell Supplement. The differentiation protocol of dopaminergic neurons from iPSCs was modified from^21^. From day 0 to day 5, the E8 complete medium was replaced with Knockout DMEM, supplemented with 15% Knockout Serum Replacement, 1x GlutaMAX, 100U/mL Penicillin-Streptomycin and 1x MEM-NEAA. The 0.1mM β-Mercaptoethanol was added freshly before changing the medium. From day 6 to day 9, the Knockout DMEM medium was gradually replaced by 25%, 50%, 75% and 100% N2 medium (DMEM/F12 with 1x N-2 supplement) respectively. The iDANs were cultured in N2 medium until day 11. From day 12 on, the iDANs were cultured in neurobasal medium supplemented with 1x B27 and 1x GlutaMAX and half medium was refreshed every two days. During the differentiation and lineage specification, different combinations of small molecules and/or growth factors were added at different time points: 10μM LDN-193189 (Stemgent Inc.) from day 1 to day 11; 10μM SB431542 (Stemgent Inc.); from day1 to day 5; 2μM Purmorphamine (Stemgent Inc.); from day 2 to day 11; 100ng/mL recombinant human fibroblast growth factor 8a (FGF-8a, R&D system) from day 2 to day 11; 3μM CHIR99021 (Stemgent Inc.); from day 3 to day 12; 100ng/ml recombinant human sonic hedgehog (C24II, SHH, R&D system); From day 12 to the end of the differentiation, 20ng/ml recombinant brain derived neurotrophic factor (BDNF, Peprotech), 20ng/ml recombinant glial-derived neurotrophic factor (GDNF, Peprotech), 0.5mM adenosine 3’,5’-cyclic monophosphate (cAMP, Enzo Life Science), 2ng/ml transforming growth factor beta 3 (TGFβ3, Millipore), 200μM ascorbid acid (AA, Sigma) and 10nM γ-secretase inhibitor IX (DAPT, Millipore, Calbiochem) were added into the medium. The cells were passaged only when they were 100% confluent using accutase. At day 20, the iDANs were split into single cells and 1-2x 10^4^/cm^2^ cells were replated on a poly-L-ornithine (Sigma, 50 μg/ml) and murine Laminin (Sigma, 10 μg/ml) double-coated plate. From day 22 on, 10μM DAPT was used to promote iDAN maturation. Rat astrocytes (prepared and plated according to^22^) were cocultured with iDANs from day 24 to further promote maturation.

### Gene expression analysis

To analyze the gene expression profile, the RNA was isolated from iPSCs and differentiated neurons with the RNeasy Mini Kit (Qiagen) according to the manufacturer’s instructions. 0,5-1ug RNA was retro-transcribed into cDNA by the iScript cDNA Synthesis Kit (Bio-Rad Laboratories, Inc) according to the manufacturer’s instructions. The gene expression of iPSCs and differentiated iDANs was measured by quantitative real-time PCR (qRT-PCR) which was performed with the Applied Biosystems 7500 Fast Real-Time PCR System using Power SYBR Green Master Mix (Applied Biosystems) and qRT-PCR primers (listed in Supplementary table S1). Beta-2-Microglobulin (B2M) was used as reference gene. The Ct value of each target gene was normalized against the Ct value of the reference gene [ΔCt= [Ct(target)-Ct(B2M)]. The relative expression was calculated as 2^−ΔΔCt^ and represented as fold change of gene expression when compared to corresponding control conditions[2^−ΔΔCt^=2^ΔCt(target)-ΔCt(control)^].

### Immunocytochemistry

The cells were fixed with 4% paraformaldehyde/4% sucrose(v/v)(Sigma) in PBS for 15min at room temperature (RT) before the immunocytochemistry experiment. The non-specific binding of antibodies was avoided by incubating the cells with 5% normal goat serum(Thermo Fisher Scientific)/ 0.4% Triton X-100(Sigma)/1% Glycine(Sigma) in PBS (blocking solution) at RT for one hour. The primary and secondary antibodies which were also diluted in the blocking solution and applied overnight at 4°C or 1h at RT. Cell nuclei were stained with Hoechst (1:10000) and the coverslips were mounted with DAKO fluoromount medium (Agilent). The primary antibodies were used: mouse anti-MAP2 (1:1000; Sigma M4403); guinea pig anti-MAP2 (1:1000; Synaptic Systems 188004); guinea pig anti-synapsin 1/2 (1:1000; Synaptic Systems 106004); mouse anti-TH (1:200; Sigma TH-16). Secondary antibodies used were: goat anti-guinea pig Alexa Fluor 568 (1:1000, Invitrogen A-11075); goat anti-rabbit Alexa Fluor 488 (1:1000, Invitrogen A-11034); goat anti-mouse Alexa Fluor 488 (1:1000, Invitrogen A-11029); goat anti-mouse Alexa Fluor 568 (1:1000, Invitrogen A-11031).

### Microelectrode array and data analysis

The neuronal network activity was measured using 6-Well Microelectrode arrays devices (Multichannel Systems, MCS GmbH, Reutlingen, Germany) consisted of 60 TiN(titanium nitride)/SiN(silicon nitride) planar round electrodes (30 μm diameter; 200 μm center-to-center interelectrode distance) divided into 6 separated wells. Each well was characterized by 9 recording electrodes, arranged in a 3 × 3 square grid, and a ground electrode. The activity of all cultures was recorded for 20 min by means of the MEA60 System (MCS GmbH, Reutlingen, Germany) with a high pass filter (Butterworth, 100Hz cutoff frequency). During the recording, the temperature was maintained at 37°C by means of a controlled thermostat (MCS GmbH, Reutlingen, Germany). Additionally, evaporation and pH changes of the medium was prevented by continuous perfusion with carbogen (95% O_2_, 5% CO_2_) inside a humidified compartment with an open connection to the MEA. After 1200x amplification, signals were sampled at 10 kHz and acquired through MC-Rack software (MCS GmbH, Reutlingen, Germany).

Data analysis was performed off-line by using a custom software developed in MATLAB (The Mathworks, Natick, MA, USA): SPYCODE ^22,52^, which collects a series of tools for processing multichannel neural recordings. Spikes and bursts were detected by using the Precise Timing Spike Detection algorithm (PTSD)^53^ and the Burst Detection algorithm^54^. The mean firing rate (spikes/s) of the network was computed by averaging the firing rate of each channel with all the active electrodes of the MEA. The burst was computed as at least 5 spikes in a burst with a maximal inter-spike-interval of 80 milliseconds. The network burst was defined as synchronized bursts that occurs in >50% of the active channels. And the network burst rate (burst/min) was calculated as the amount of network bursts per minute.

### Single-cell electrophysiology

Experiments were conducted on DIV55 iDANs. The coverslip with neurons was transferred the to a recording chamber continuously perfused with oxygenated (95% O_2_ / 5% CO_2_) and heated (32°C) recording ACSF containing (in mM): 124 NaCl, 3 KCl, 1.25 NaH_2_PO_4_, 2 CaCl_2_, 1 MgCl_2_, 26 NaHCO_3_, 10 Glucose. Patch pipettes (5.5-7.5 MΩ) were made from borosilicate glass capillaries and filled with intracellular recording solution containing (in mM): 130 K-Gluconate, 5 KCl, 10 HEPES, 2.5 MgCl_2_, 4 Na_2_-ATP, 0.4 Na_3_-ATP, 10 Na-phosphocreatine, 0.6 EGTA (adjusted to pH 7.25 and osmolarity 290 mosmol). Activity was recorded using a Digidata 1440A digitizer and a Multiclamp 700B amplifier (Molecular Devices). Sampling rate was set at 20KHz and a lowpass 1KHz filter was used during recording. Series resistance was monitored on-line and cells were discarded if series resistance increased above 1:10 of membrane resistance.

#### Spontaneous postsynaptic currents

sEPSCs were recorded in drug free recording medium at a holding potential of −60mV.

#### Intrinsic properties

Intrinsic properties were recorded in drug free medium. Current-voltage responses were recorded by 1 second stepwise injection of current in 5 pA increments. Current injection continued until clear action potentials were visible or until a maximum current of 100 pA was injected. Intrinsic properties were calculated off-line using Clampfit 10.7 software (Molecular Devices, San Jose, CA). Statistical comparison was conducted in PRISM (Graphpad PRISM 7.00, Graphpad Software, San Diego, CA).

### Neuronal morphology reconstruction and data analysis

Wide field fluorescent images of the fixed and MAP2-labelled iPSC derived dopaminergic neurons were taken at 20x magnification using ApoTome microscopy (Zeiss Axio Imager Z1/Z2). The images were stitched using Fiji 2017 software with the stitching plugin and digitally reconstructed using Neurolucida 360 (Version 2017.01.4, Microbrightfield Bioscience, Williston, USA) to create an overlay drawing of the somatodendritic morphology. Only the neurons that have at least two primary dendrites and at least one dendritic branch point were selected for the reconstruction.

The quantitative analysis of the somatodendritic organization of the dopaminergic neurons was conducted using Neurolucida 360 Explorer (Version 2017.02.7, Microbrightfield Bioscience, Williston, USA). For the somatodendritic analysis, the soma area, number of dendritic nodes, dendritic length, quantity of primary dendrites, as well as the covered surface by convex hull 2D analysis were determined. Furthermore, Sholl analysis was performed to investigate the dendritic complexity^55^. The dendritic length of the neurons was measured within a series of concentric circles at 25 μm interval from the soma. All the morphological data were acquired and analyzed blindly to the genotype of the neurons. For the somatodendritic properties and Sholl analysis, significance was determined using IBM SPSS Statistics (version 24.0, IBM, Armonk, USA) to run Wilks’ Lambda multivariate analysis of variance (MANOVA) followed by the Bonferroni post-hoc.

### Statistical analysis

Statistical analysis of the data was performed with GraphPad Prism 5.0.3 (GraphPad Software, Inc, USA). And the distribution of data is expressed as mean ± SEM. Mann-Whitney U test, unpaired Student’s T test or one-way ANOVA with Tukey’s correction for multiple-comparisons was used for statistical analysis. p < 0.05 was considered significant.

## Supplementary material

**Figure S1:**
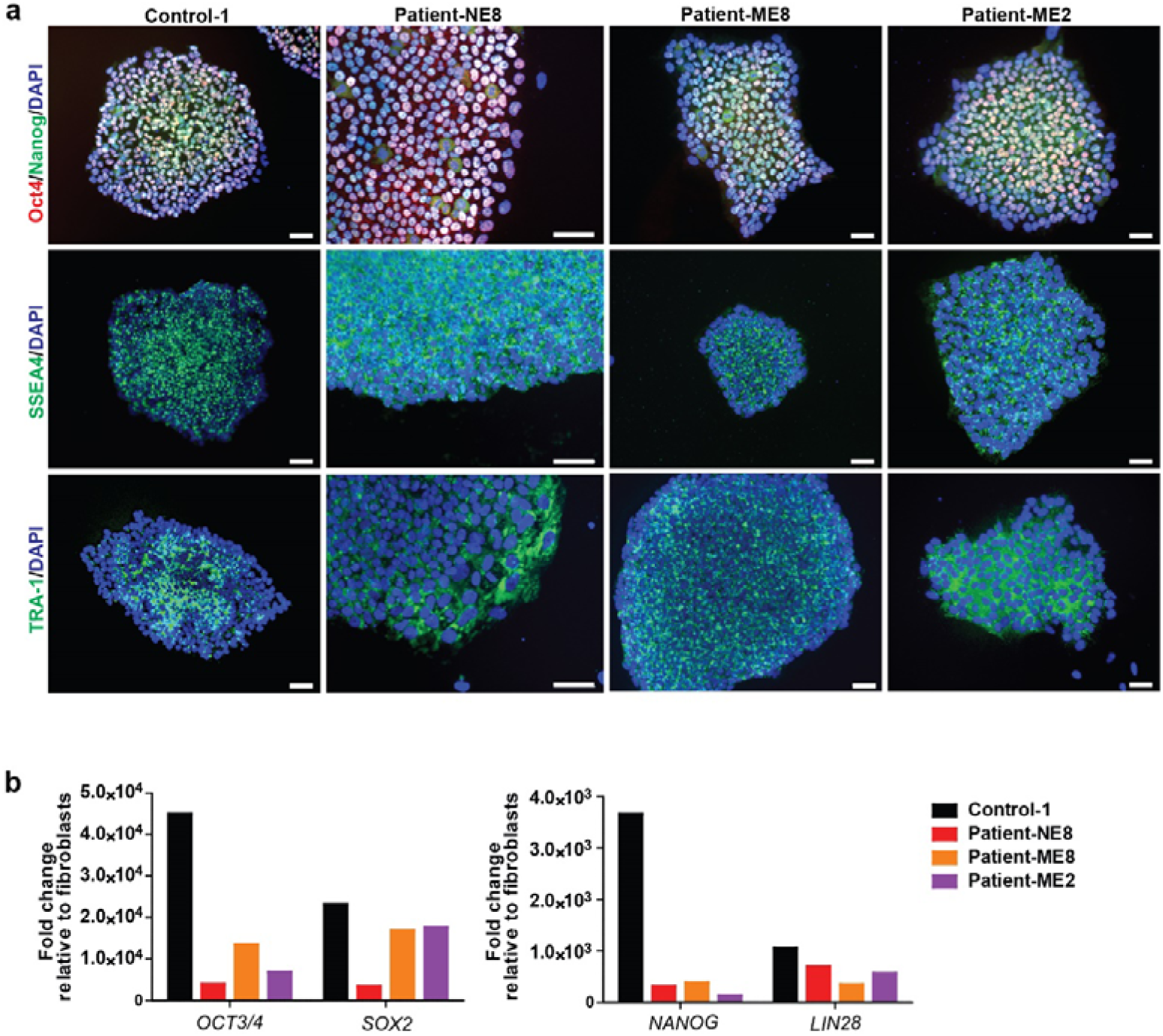
Characterization of iPSC lines with immunocytochemistry and qPCR. (a) All the lines were positive for the staining of pluripotent makers Oct 4, Nanog, SSEA4 and TRA-1. (b) The expression of pluripotent markers increased 10^2^ to 10^5^-folds compared to the average expression of corresponding fibroblasts (Scale bar=50 μm).

**Figure S2:**
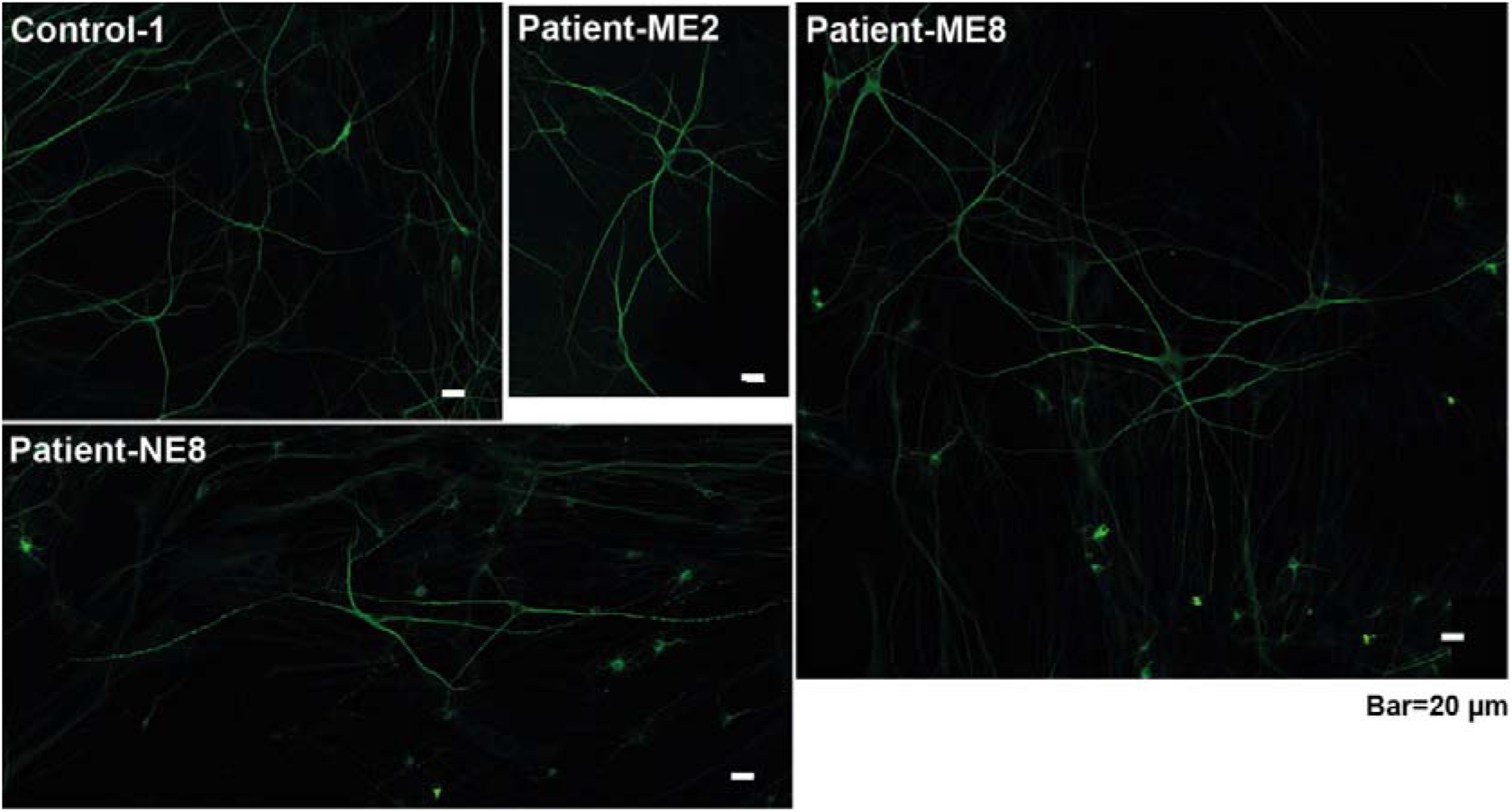
Representative MAP2 staining for reconstruction and Sholl analysis of neuron dendritic length. Representative MAP2 immunostaining of iDANs for the reconstruction of day 73 (Scale bar=20 μm).

**Figure S3:**
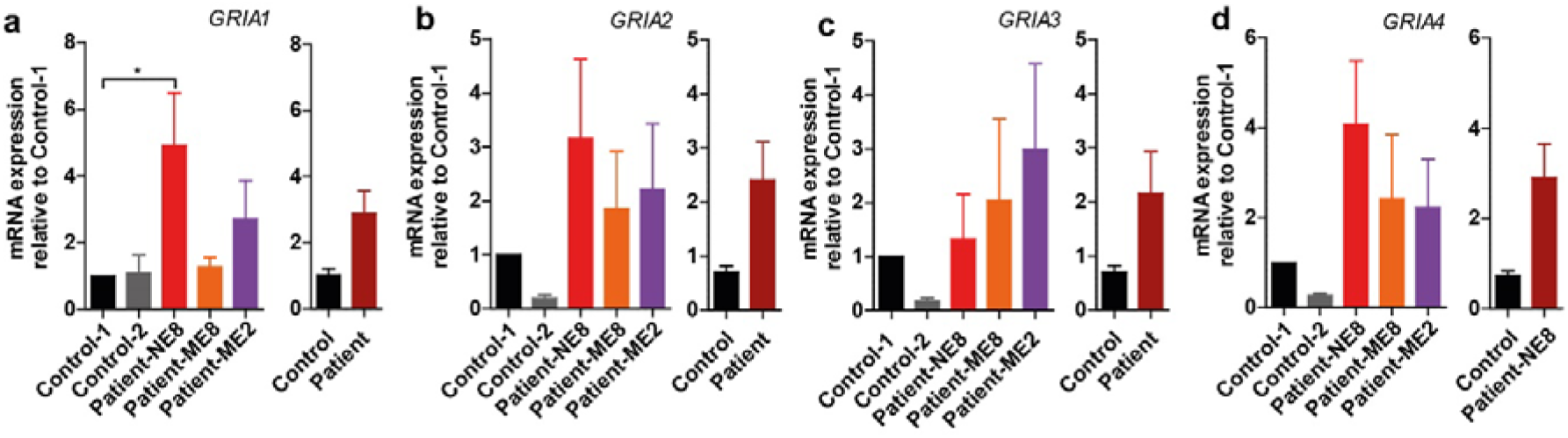
No significant difference of the expression of AMPA receptor subunits between control and patient. The expression of genes encoding AMPA receptor subunits *GRIA1*(a), *GRIA2*(b), *GRIA3*(c) and *GRIA4*(d) were analyzed by qPCR in day 73 iDANs. The mRNA expression were normalized to the mean of Control-1. Statistical analysis was done with Mann-Whitney U-test between the group of control and patient and one way ANOVA followed by Tukey’s multiple comparison test for different genotypes. Data are mean ±SEM, asterisks indicate significant differences between genotypes. *p<0.05. (n≥ 5).

**Figure S4:**
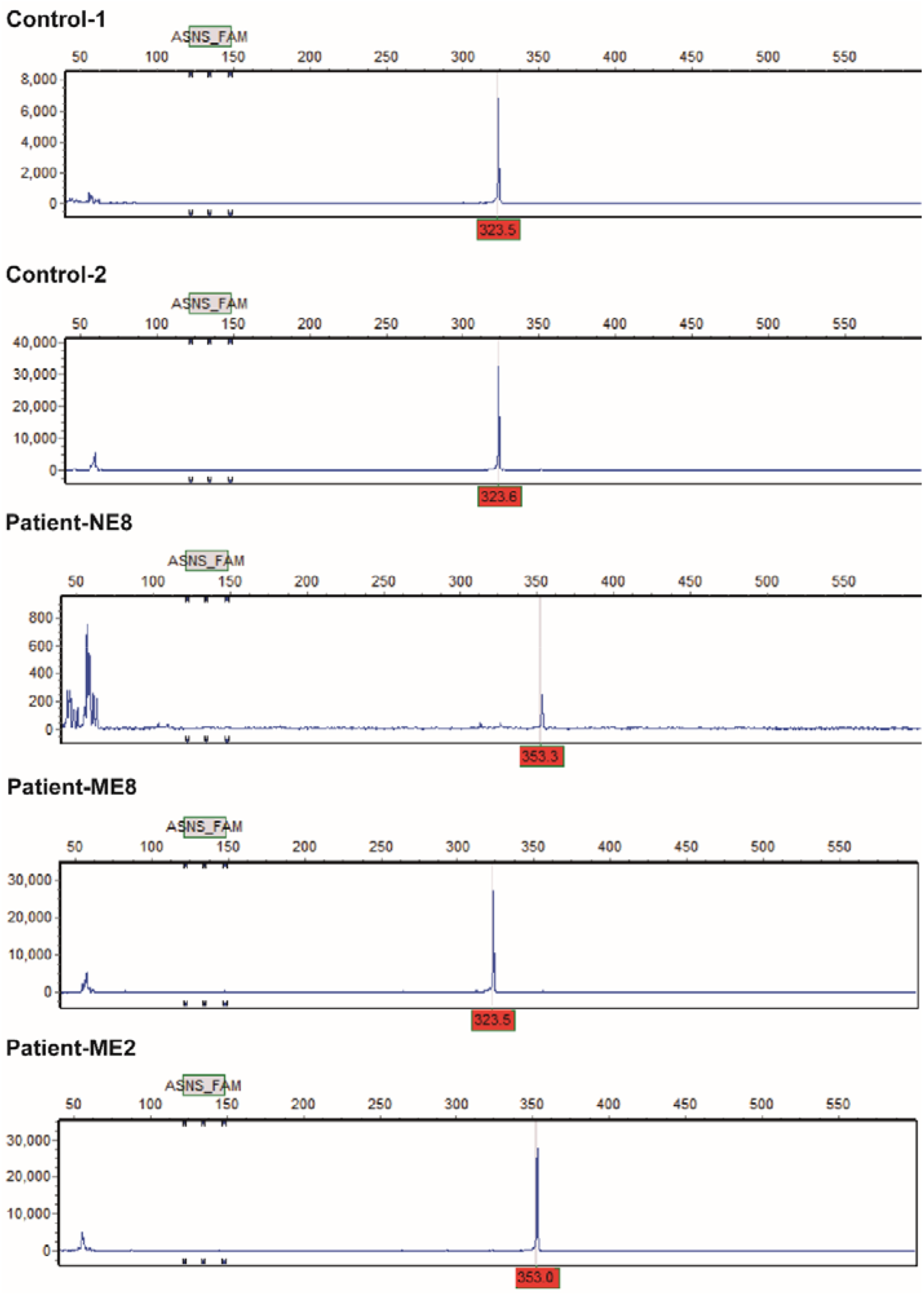
Variable number tandem repeats (VNTRs) polymorphism of *MAOA* promoter.(Corresponding to table S2). The peaks at 323 position indicate the individuals have a 3R allele and the peaks at 353 position indicate the individuals carry a 4R allele.

**Supplementary table S1:**
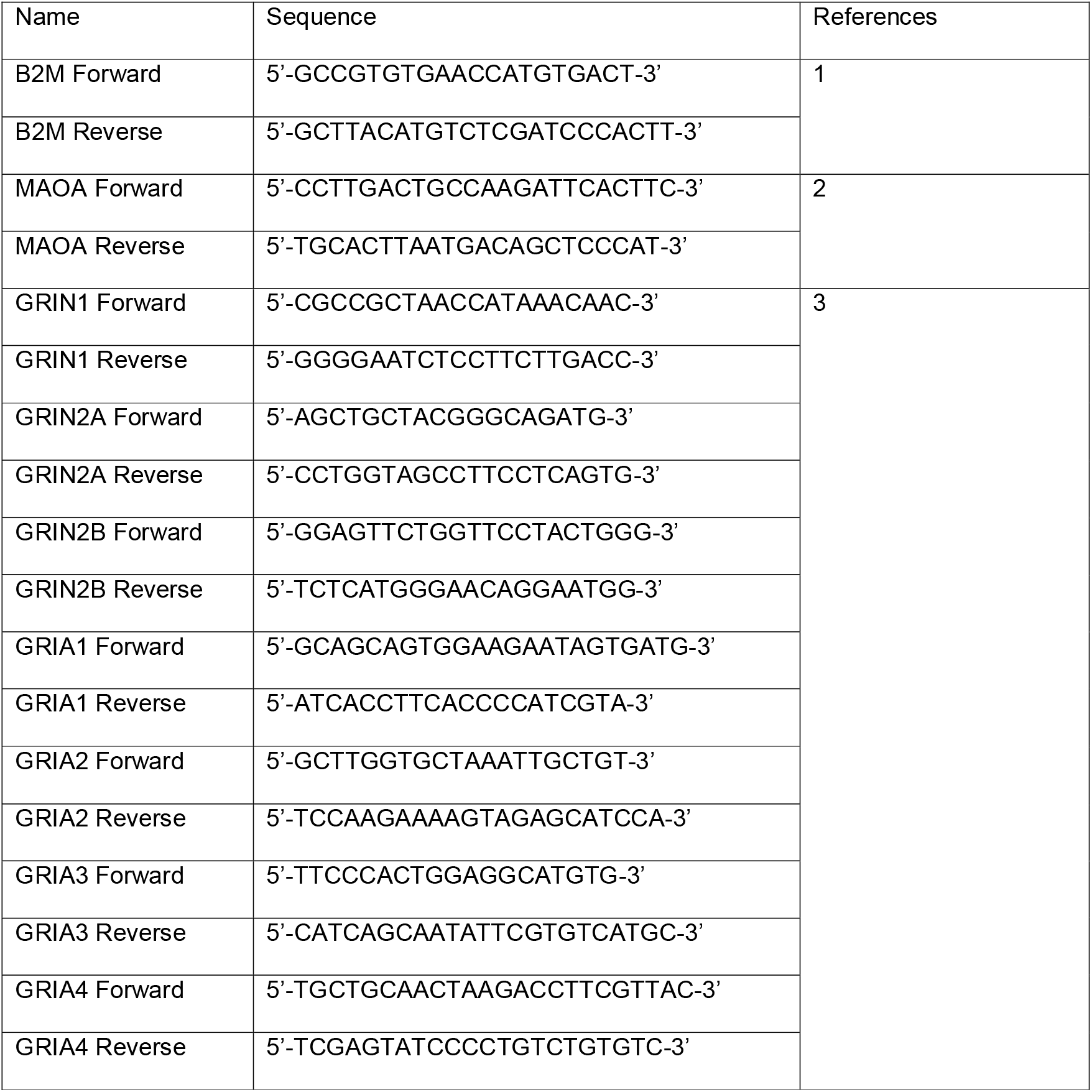
primers for qRT-PCR

**Supplementary table S2:**
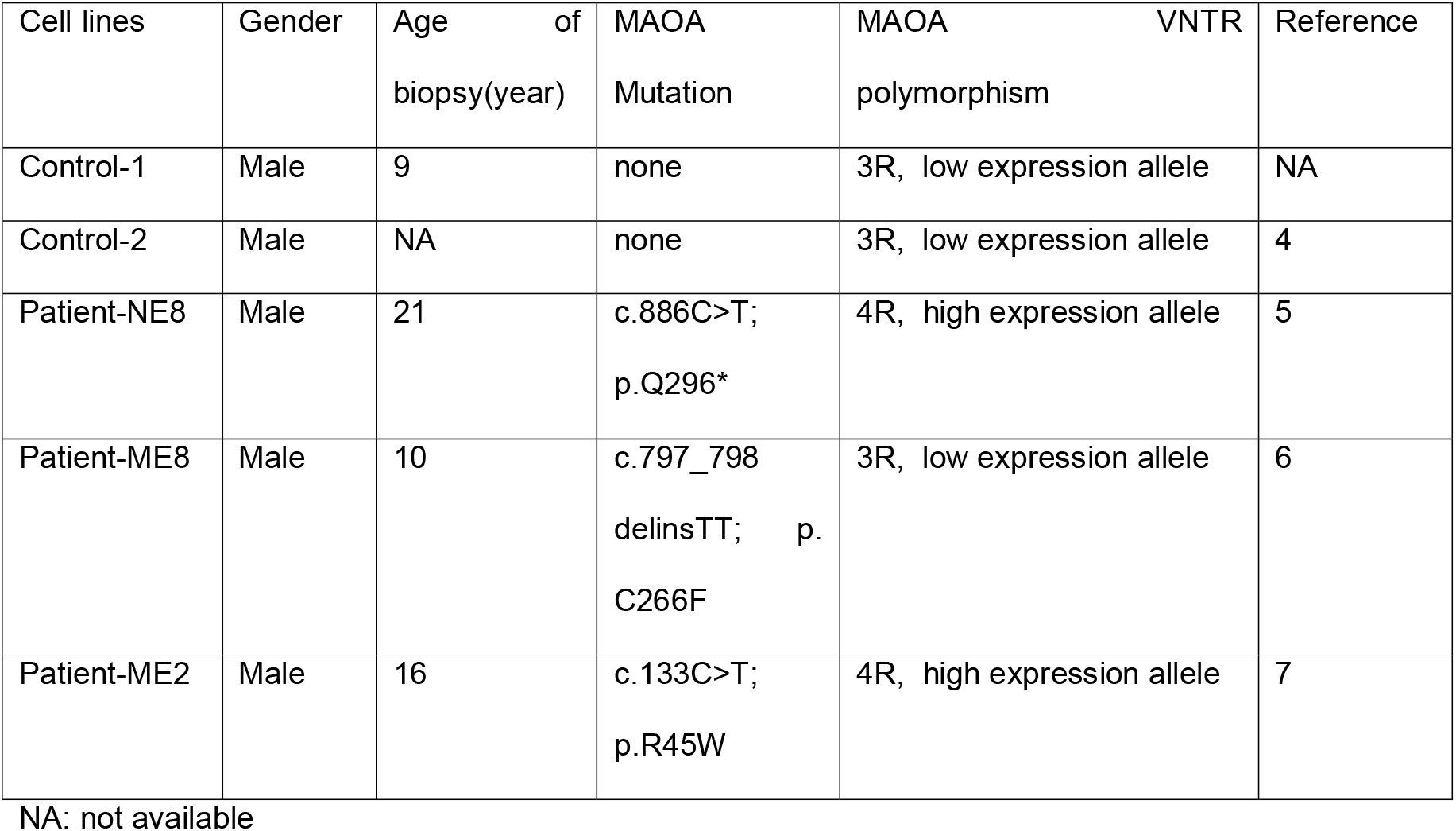
information about the cell lines used in the study

| Cell lines | Gender | Age of biopsy(year) | MAOA Mutation | MAOA VNTR polymorphism | Reference |
| --- | --- | --- | --- | --- | --- |
| Control-1 | Male | 9 | none | 3R, low expression allele | NA |
| Control-2 | Male | NA | none | 3R, low expression allele | 4 |
| Patient-NE8 | Male | 21 | c.886C>T;<br>p.Q296* | 4R, high expression allele | 5 |
| Patient-ME8 | Male | 10 | c.797_798<br>delinsTT; p.<br>C266F | 3R, low expression allele | 6 |
| Patient-ME2 | Male | 16 | c.133C>T;<br>p.R45W | 4R, high expression allele | 7 |

NA: not available

## Supplementary methods

Genotyping of the MAOA promoter VNTR polymorphism: The genomic DNA was amplified with 1x AmpliTaq Gold® 360 Master Mix (Life Technologies) and 0.33 mM of fluorescently labeled forward primer (FAM-5’-ACAGCCTGACCGTGGAGAAG-3′) and reverse primer (5′-GAACGGACGCTCCATTCGGA-3′) in a total volume of 7,5μl using the protocol: 95°C for 10 min followed by 35 cycles of denaturation for 30 s at 95°C, 30 s annealing at 60°C, primer extension at 72°C for 1min and a final extension at 72°C for 10 min. Fragment length analysis of the PCR product was performed by an automated capillary sequencer ABI3730 (Applied Biosystems, Nieuwerkerk a/d Ijssel, The Netherlands) using standard conditions (1 ul of the 1:20 diluted PCR product together with 9.7 ul formamide and 0.3ul GeneScan-600 Liz Size Standard TM (Applied Biosystems, Nieuwekerk aan den Ijsel, the Netherlands). Results were analyzed with GeneMarker version 2.6.7 (SoftGenetics, US).

